# NGSphy: phylogenomic simulation of next-generation sequencing data

**DOI:** 10.1101/197715

**Authors:** Merly Escalona, Sara Rocha, David Posada

## Abstract

**Motivation:** Advances in sequencing technologies have made it feasible to obtain massive datasets for phylogenomic inference, often consisting of large numbers of loci from multiple species and individuals. The phylogenomic analysis of next-generation sequencing (NGS) data implies a complex computational pipeline where multiple technical and methodological decisions are necessary that can influence the final tree obtained, like those related to coverage, assembly, mapping, variant calling and/or phasing.

**Results:** To assess the influence of these variables we introduce NGSphy, an open-source tool for the simulation of Illumina reads/read counts obtained from haploid/diploid individual genomes with thousands of independent gene families evolving under a common species tree. In order to resemble real NGS experiments, NGSphy includes multiple options to model sequencing coverage (depth) heterogeneity across species, individuals and loci, including off-target or uncaptured loci. For comprehensive simulations covering multiple evolutionary scenarios, parameter values for the different replicates can be sampled from user-defined statistical distributions.

**Availability:** Source code, full documentation and tutorials including a quick start guide are available at http://github.com/merlyescalona/ngsphy.

## 1. Introduction

Next-generation sequencing (NGS) technologies facilitate nowadays the obtention of large “phylogenomic” datasets with hundreds or thousands of loci with several individuals from multiple species (McCormack *et al.*, 2013). At the same time, genome-wide data have brought a renewed interest in the discordance between gene trees and species trees and in methods of simulation and phylogenetic reconstruction that deal with large amounts of loci and potential sources of phylogenetic incongruence (Mallo and Posada, 2016). Importantly, the assembly of multiple sequence alignments from NGS reads is not free from errors and biases. There are many variables that might interfere with the accuracy of the final gene trees and species trees inferred from NGS data, from experimental-design aspects such as number of samples, sequencing technology or depth of coverage (herein coverage), to methodological decisions along the processes of assembly or mapping, orthology inference, variant and genotype calling, or phasing. In this context, the simulation of NGS reads can be quite useful to properly understand and improve the NGS phylogenomic pipeline. However, at the phylogenomic scale we need to be able to simulate NGS data from multiple gene families or loci with potentially discordant phylogenies, and represented by multiple individuals. To the best of our knowledge, the only software which generates NGS data from phylogenies is TreeToReads (McTavish *et al.*, 2017). Although this is a very useful tool for the simulation of NGS data along a single gene tree, it cannot directly simulate data along multiple gene trees, diploid individuals, read counts, or consider coverage heterogeneity across individuals and loci, being therefore not easily applicable to the species tree / phylogenomic scenario. In order to solve this gap, here we introduce NGSphy, an easy-to-use pipeline to generate NGS data (read counts or Illumina reads) from multiple loci belonging to multiple haploid/diploid individuals and species under the gene tree / species tree paradigm, and with different options to control coverage variation across species, loci and individuals.

## 2. Description and implementation

NGSphy is an open-source tool written in Python that takes advantage of the NumPy (van der Walt *et al.*, 2011) and Dendropy (Sukumaran and Holder, 2010) libraries. Its workflow is depicted in Supplementary Figure S1. Parameter values and options for the simulations are specified in a settings file. Arguments and conditions for the different replicates can be sampled from user-defined statistical distributions, or set as fixed. In the simplest scenario, the user just needs to specify a single gene tree, a substitution model, and the sequencing design. Optionally, the user can provide an ancestral or tip nucleotide sequence that will be used at the root of the tree to start the simulation. Otherwise, the root sequence is simulated according to the stationary frequencies of the specified substitution model. Nucleotide sequence alignments are evolved using INDELible (Fletcher and Yang, 2009) or a version of it modified by us in order to force the sequence at the root (see online documentation). For more complex scenarios, NGSphy is able to read directly the output of SimPhy (Mallo *et al.* 2015), a program that simulates multiple gene trees evolving within a species tree under incomplete lineage sorting, gene duplication and loss, horizontal gene transfer or gene conversion, plus the corresponding multilocus alignments obtained with INDELible. Before producing the NGS reads, the genomic sequences at the tip of each gene tree are assigned to haploid (directly) or diploid (by random sampling within species) individuals.

With the trees, multiple sequence alignments and individuals in place, NGSphy can directly produce read counts for single-nucleotide variants (SNVs) or generate Illumina reads using ART (Huang *et al.*, 2012). Read counts are produced taking into account a user-defined error rate. The sequencing In this case, coverage (depth) at each position is sampled from a Negative Binomial distribution whose mean is the sampled coverage for the corresponding species, locus and individual. For diploid individuals, coverage is further split among homologous chromosomes with equal probability. If desired, genotype likelihoods for every site can be computed as in GATK (Mckenna *et al.*, 2010). The simulated read counts are written to a set of VCF files, one per locus. In addition, NGSphy can call ART with the parameter values indicated in the settings file and produce Illumina reads for each individual in ALN, BAM and/or FASTQ format. ART is a very flexible program that uses customized read error model parameters and quality profiles.

Importantly, NGSphy can simulate the variation of coverage that may occur in NGS due to differences in quantity or quality of DNA samples, technical problems when generating libraries or genomic changes in GC content. Coverage variation is implemented hierarchically. For each replicate, an experiment-wide coverage is assigned according to the user–fixed or sampled from a distribution of choice. Coverage variation among loci and individuals is introduced by sampling multipliers from user-defined Gamma distributions, while coverage variation across species/taxa can be directly specified (see online documentation for details). For targeted-sequencing experiments NGSphy can simulate off-target loci (not targeted but captured with reduced coverage), uncaptured loci, or a coverage decay related to the phylogenetic distance to a selected reference sequence (Bragg *et al.*, 2016). Finally, NGSphy allows for multi-threaded, parallel runs. In addition, it can generate job templates for execution in computational clusters.

## 3. Validation and case study

To validate NGSphy we performed different experiments and sanity checks, inspecting multiple runs and output files. We carried out a small test where the generating gene tree was always properly recovered from the simulated alignments (Supplementary data: validation test). We also included a small case study where the number of SNP calls from NGS reads grows rapidly, as expected, with sequencing coverage (Supplementary data: use case).

## Acknowledgements

We thank Diego Mallo, the Phylogenomics lab (http://darwin.uvigo.es), Rute R. da Fonseca and Anders Albrechtsen for their feedback.

## Funding

This work was supported by the Spanish Government (research grants BFU2012-33038 and BFU2015-63774-P to D.P.; FPI graduate fellowship BES-2013-067181 to M.E. and Juan de la Cierva postdoctoral fellowship FPDI-2013-17503 to S.R.) and the European Research Council (ERC-617457-PHYLOCANCER to D.P.)

## Conflict of Interest

none declared.

